# Genetic tracing of market wildlife and viruses at the epicenter of the COVID-19 pandemic

**DOI:** 10.1101/2023.09.13.557637

**Authors:** Alexander Crits-Christoph, Joshua I. Levy, Jonathan E. Pekar, Stephen A. Goldstein, Reema Singh, Zach Hensel, Karthik Gangavarapu, Matthew B. Rogers, Niema Moshiri, Robert F. Garry, Edward C. Holmes, Marion P. G. Koopmans, Philippe Lemey, Saskia Popescu, Andrew Rambaut, David L. Robertson, Marc A. Suchard, Joel O. Wertheim, Angela L. Rasmussen, Kristian G. Andersen, Michael Worobey, Florence Débarre

## Abstract

Zoonotic spillovers of viruses have occurred through the animal trade worldwide. The start of the COVID-19 pandemic was traced epidemiologically to the Huanan Wholesale Seafood Market, the site with the most reported wildlife vendors in the city of Wuhan, China. Here, we analyze publicly available qPCR and sequencing data from environmental samples collected in the Huanan market in early 2020. We demonstrate that the SARS-CoV-2 genetic diversity linked to this market is consistent with market emergence, and find increased SARS-CoV-2 positivity near and within a particular wildlife stall. We identify wildlife DNA in all SARS-CoV-2 positive samples from this stall. This includes species such as civets, bamboo rats, porcupines, hedgehogs, and one species, raccoon dogs, known to be capable of SARS-CoV-2 transmission. We also detect other animal viruses that infect raccoon dogs, civets, and bamboo rats. Combining metagenomic and phylogenetic approaches, we recover genotypes of market animals and compare them to those from other markets. This analysis provides the genetic basis for a short list of potential intermediate hosts of SARS-CoV-2 to prioritize for retrospective serological testing and viral sampling.

## Introduction

Many of the earliest known cases of COVID-19 worked at or visited the Huanan Seafood Wholesale Market (“Huanan market”) in the city of Wuhan, a link first made by clinicians at different hospitals throughout the city (Wuhan Municipal Health Commission 2019; Worobey 2021; Worobey et al. 2022). Retrospective review of early COVID-19 cases identified 174 patients with onset in December 2019, 32% of whom had an ascertained link to this location within a city of about 12 million (World Health Organization 2021). While initial case finding could have preferentially identified market linked cases, a geospatial analysis of residences of the cases with no identified link to the Huanan market showed that they lived unexpectedly close to and centered around the market (Worobey et al. 2022), even though geographic proximity was not used as a case criterion (Chen et al. 2020; Li et al. 2020; Wang et al. 2020; World Health Organization 2021). Additionally, excess pneumonia deaths were first reported in the city districts surrounding the Huanan market (World Health Organization 2021; Holmes et al. 2021), and retrospective serosurveys of Wuhan confirmed that a larger proportion of residents contracted COVID-19 in these districts (Li et al. 2021; He et al. 2021).

The genomic epidemiology of SARS-CoV-2 shows that there were very few human infections before the earliest reported market case with onset on December 10^th^, 2019 (Pekar et al. 2022). The time of the most recent common ancestor (tMRCA) is estimated to be late November to December 2019 (Duchene et al. 2020; Lu et al. 2020; Giovanetti et al. 2020; Gómez-Carballa et al. 2020; Li et al. 2020; Pekar et al. 2021), and the estimated median timing of the primary case mid to late November (Pekar et al. 2022; Jijón et al. 2023). A phylodynamic analysis of the epidemic’s size by December 1 estimated it to be between 1–83 infections and 0–2 hospitalizations (95% highest posterior density intervals) (Pekar et al. 2022). These estimates are consistent with surveillance and retrospective testing that have found no evidence of substantial community transmission of COVID-19 prior to December 2019 (Chang et al. 2023, 2021; Kong et al. 2020; Tao et al. 2020; World Health Organization 2021).

Early SARS-CoV-2 sequences belong to two lineages, denoted A and B, separated by two characteristic genome mutations (C8782T and T28144C). While the rooting of SARS-CoV-2 between these two haplotypes is uncertain (Pekar et al. 2022), the initial observation that SARS-CoV-2 genomes from cases with direct market contact were lineage B led to the proposal that the market was an amplification event that occurred after lineage A community transmission unrelated to the market (Bloom 2021). The geographic proximity of two early lineage A cases to the market, however, instead suggested that the lineage was present (Worobey et al. 2022), and this was further confirmed when lineage A was identified in an environmental sample from the Huanan market (Liu et al. 2023). The linkage of both lineages to the market is consistent with phylogenetic evidence of at least two sustained zoonotic spillovers of SARS-CoV-2 into humans (Pekar et al. 2022). The high intensity of contact between humans and animals in markets (Pruvot et al. 2019) suggests that once animals infected with a highly transmissible virus arrive in a market, multiple zoonotic events are primed to occur.

In February 2020, China’s government enacted a far-reaching ban on the sale of wildlife for human consumption (Koh, Li, and Lee 2021). A similar decision had followed the second emergence of SARS-CoV-1 in winter 2003–2004 (Shi and Hu 2008); both were intended to limit the further spread of either virus within the animal trade. As with SARS-CoV-2, SARS-CoV-1 was first detected over 1,000 kilometers from its closest identified bat virus relatives in Yunnan province and was epidemiologically linked to the wildlife trade (Xu et al. 2004). In the months after declaring the SARS outbreak, closely related viruses were found in masked palm civets (*Paguma larvata)*, common raccoon dogs (*Nyctereutes procyonoides*), and a ferret badger (*Melogale moschata*) at still-open wet markets(Guan et al. 2003), although animals from several markets and farms tested negative for SARS-CoV-1 (Shi and Hu 2008; Tu et al. 2004; Kan et al. 2005). Farmed civets in Hubei province also tested positive, indicating spread of SARS-CoV-1 among animals to the province where SARS-CoV-2 later emerged (Shi and Hu 2008).

Zoonotic spillovers in wildlife markets have long been known to present risks for viral emergence (Keusch et al. 2022). Cross-species transmissions of bat coronaviruses to mammals in the wildlife trade have also been documented among Malayan porcupines and hoary bamboo rats (Huong et al. 2020). Coronaviruses have been reported in masked palm civets, raccoon dogs, and Amur hedgehogs in wildlife markets (He et al. 2022), and while the closest relatives of SARS-CoV-2 to date are in bats (Temmam et al. 2022), other closely related viruses have been been found in illegally traded pangolins in Asia (Lam et al. 2020; Nga et al. 2022; Xiao et al. 2020; Wacharapluesadee et al. 2021). In rural Myanmar, individuals with wildlife exposure had higher sarbecovirus seropositivity, possibly indicating rare spillover to humans as well (Evans et al. 2023).

The Huanan market was the location with the most wildlife vendors in Wuhan, a city of over 12 million people with four sustained live animal markets (Xiao et al. 2021). Several vendors were documented to be illegally selling live animals such as raccoon dogs, civets, bamboo rats (*Rhizomys pruinosus* and/or *Rhizomys sinensis*), Malayan porcupines (*Hystrix brachyura*), Amur hedgehogs (*Erinaceus amurensis*), and Asian badgers (*Meles leucurus*) in late fall of 2019 (Xiao et al. 2021; Worobey et al. 2022). Most wildlife vendors in the market were located in the west wing, which was also where the earliest and the majority of market COVID-19 cases worked (Worobey et al. 2022). In early 2020, Liu et al. collected environmental samples from the Huanan market. These samples were analyzed by SARS-CoV-2 reverse-transcription quantitative PCR (qPCR) and metatranscriptomic next-generation sequencing (mNGS) (Liu et al. 2023). In addition to environmental sampling, Liu et al. performed qPCR testing of some mammalian wildlife at the market, but this was limited to species now known as unlikely intermediate hosts of SARS-CoV-2 such as stray weasels, rats, cats, and dogs, as well as carcasses of one sheep, two wild boars, six bamboo rats, six badgers, six muntjacs, 16 hedgehogs, and 52 rabbits (Liu et al. 2023). No samples from raccoon dogs, civets, or porcupines on sale in the market were tested by qPCR, and no serology from animals in the market has been described. However, Liu et al. reported the genetic detection in environmental samples of several animal genera of potential interest (Liu et al. 2023).

Here, we analyze the data from the market generated and shared by Liu et al. using multiple novel genomic approaches. We first demonstrate that SARS-CoV-2 genetic diversity within the Huanan market reflects the tMRCA of the global pandemic, a finding consistent with its emergence within the market. We characterize the genetic material from mammals present in market metatranscriptomes at the species level, and thereby document their presence in samples and stalls with SARS-CoV-2. We further identify multiple additional viruses most likely shed by live mammals sold at the market. Finally, we reconstruct mitochondrial genotypes of putative intermediate hosts in the market for identification of subspecies and their putative geographic origins. Taken together, these new analyses provide a new and precise picture of the genetic signature of wildlife mammals, their viruses, and SARS-CoV-2 that were present at the Huanan market as the COVID-19 pandemic began.

## Results

### SARS-CoV-2 genetic diversity linked to the Huanan market is consistent with market emergence

If the Huanan market was the origin of the viral transmission chain that led to the COVID-19 pandemic, then the common ancestor of market-associated viral genotypes should be equivalent to the common ancestor of the pandemic, given sufficient sampling. To test this hypothesis, we first assess intra-sample variation of the SARS-CoV-2 environmental genomes from the Huanan market and confirm that one sample (A20) conclusively contained lineage A. We then perform phylodynamic inference to compare the genetic diversity of SARS-CoV-2 within the market to its genetic diversity globally, and found that the tMRCA of market-associated genomes reflects the global tMRCA of the pandemic (**Figure 1A**).

**Figure 1:**
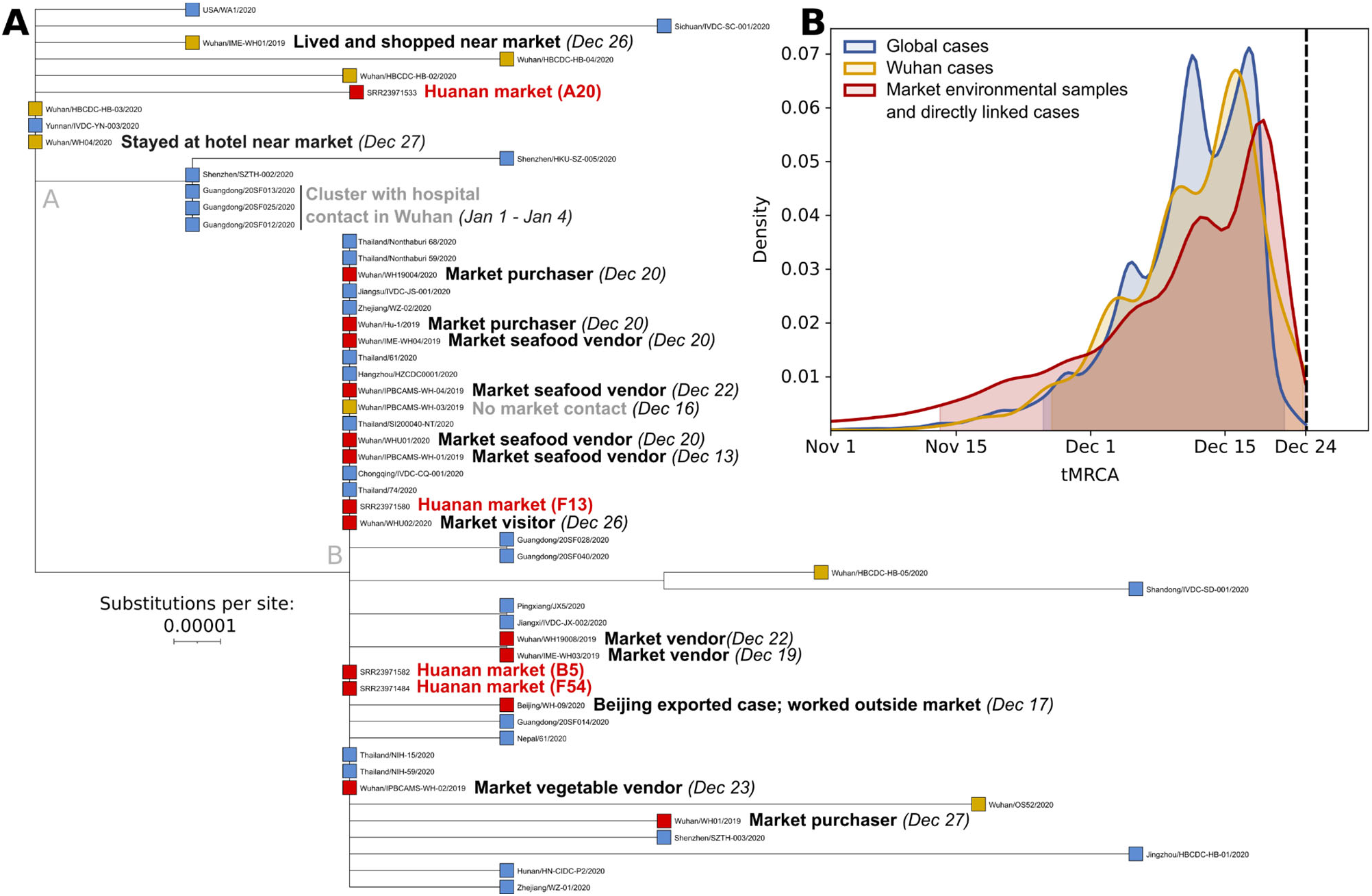
Phylogenetic analysis of SARS-CoV-2 from early COVID-19 cases and virus sequences obtained from the Huanan market. (**A**) Phylogenetic tree of SARS-CoV-2 viral genomes collected before Jan 20, 2020. Tip colors correspond to different samples (red: market environmental samples and directly linked cases; yellow: Wuhan cases, with those indirectly linked to the Huanan market labeled; blue: all global cases). Symptom onset dates for cases are shown when known. The branch leading to A20 is slightly longer than other sequences with two substitutions from the root due to its slightly greater proportion (2.2%) of undetermined nucleotides. (**B**) tMRCA estimates for SARS-CoV-2 viral sequences from samples collected by February 14^th^ 2020, obtained from globally collected viral samples (blue), samples obtained from Wuhan (yellow), and samples from the Huanan market or cases with direct market contact (red). The 95% highest posterior density (HPD) interval of each distribution is highlighted and the dashed line denotes the sampling date of the earliest genome (December 24^th^ 2019).

Four near-complete SARS-CoV-2 genome sequences were recovered from environmental samples collected on January 1^st^ 2020 (Liu et al. 2023). The samples, labeled A20, B5, F13, and F54, had all been collected in the west wing of the Huanan market where most wildlife vendors were located (World Health Organization 2021). We first confirmed that the sequence from sample A20 is lineage A, whereas samples F54, F13, and B5 are lineage B, as previously reported (Liu et al. 2023). The sampled reads strongly support the A or B consensus genotype at the lineage-defining sites, indicating that the samples are neither mixtures of lineage A and B, nor intermediates between the two. The A20 consensus genome has an additional two mutations, G26262T and C6145T, separating it from the lineage A root (**Figure 1A**), but with the 6145C reference allele present at 23% frequency, supporting the presence of a haplotype 1 private mutation diverged from the MRCA of lineage A (**Supplementary Table S1**). While the exact timing of the shedding of virus present in this lineage A sample remains uncertain, a map shared during the WHO-China origins tracing joint mission had noted a case in the stall it originated from (“Origins of Covid” BMJ Webinar 2021), with onset on or before December 15^th^**(Figure S1B)**. This case was not included on the final map provided in the WHO mission report, but the report recommended follow up mapping and review of potential early clinical cases (World Health Organization 2021).

To compare the genetic diversity of SARS-CoV-2 sampled within the market to SARS-CoV-2 genetic diversity globally from early 2020 infections, we performed phylodynamic inference using BEAST (Suchard et al. 2018) (**Figure 1A**). Using the sample collection date of January 1^st^ for environmental genomes, we inferred the tMRCA of SARS-CoV-2 using sequences from three increasingly large geographical areas: cases and environmental sequences directly linked with the Huanan market (n=17) (**Supplementary Table S2**), all sequences from Wuhan (n=93), and all global sequences (n=789) from samples collected on or before February 14^th^, 2020 (**Figure 1B**). As expected due to the presence of both lineages A and B at the Huanan market, all three tMRCA distributions overlapped (their 95% highest posterior density (HPD) intervals range from Nov 13 to Dec 23), establishing that the timing of the origin of the market outbreak was indistinguishable from the timing of the origin of the global pandemic. This is consistent with previously published results that identify the ancestral MRCA of SARS-CoV-2 as either the A or B haplotypes or an unobserved intermediate between the two (Pekar et al. 2022). The association of SARS-CoV-2 lineages A and B with the Huanan market confirms that the COVID-19 outbreak within the market was not a lineage B superspreading event. Rather, the presence of both lineages A and B at the market, and the spatial association of early lineage A cases with the market (Worobey et al. 2022), are results predicted under the hypothesis that SARS-CoV-2 first emerged in the human population at the Huanan market.

### High SARS-CoV-2 positivity in and near a wildlife stall in the Huanan market

The Huanan market was sampled on multiple dates at the start of 2020, with different sampling trips having different purposes (Liu et al. 2023). On January 1^st^, the market was sampled widely with an emphasis on stalls associated with human cases: 515 samples were tested, 27 were qPCR-positive, and 25 of these SARS-CoV-2 positive samples were subsequently sequenced with metatranscriptomics. On January 12^th^, 10 samples per stall were taken from seven wildlife stalls: three were positive by qPCR for SARS-CoV-2, and this time all 70 samples were sequenced. Additional samples were collected from drains, sewage, stalls and warehouses after these two first dates until March 2020 (**Supplementary Table S3-S4**).

To determine whether the SARS-CoV-2 positivity was associated with specific stalls in the Huanan market, we conducted a spatial relative risk analysis of SARS-CoV-2 qPCR-positive samples from the January 1^st^ and 12^th^ collections, comparing the distribution of the qPCR-positive to qPCR-negative samples. The rate of qPCR-positivity was unevenly distributed within the Huanan market, with increased positivity in the southwest section (**Figure 2A**). Several clustered stalls in this section had a higher positivity rate than the average stall sampled in the market (**Figure S1**). One stall, wildlife stall A, stood out with a 30% qPCR-positive rate (three of its ten samples): a cart, a hair/feather removal machine, and a floor sample were qPCR-positive for SARS-CoV-2. Six of the 70 January 12^th^ sequenced samples contained SARS-CoV-2 sequence reads, which included the three qPCR-positive samples from wildlife stall A, two qPCR-negative samples also from wildlife stall A, and one sample from the interior of a freezer in nearby wildlife stall B (**Supplementary Table S5**; **Figure 2B**). Both qPCR testing and mNGS, therefore, independently identify SARS-CoV-2 RNA in and around wildlife stall A (**Figure 2A, 2B**). Although SARS-CoV-2 read counts are low, this is consistent with precedent for untargeted environmental sequencing of SARS-CoV-2, in which viral RNA can be nearly undetectable even in PCR-positive samples where the overwhelming majority of sequences are microbial (Rothman et al. 2021; Crits-Christoph et al. 2021).

**Figure 2:**
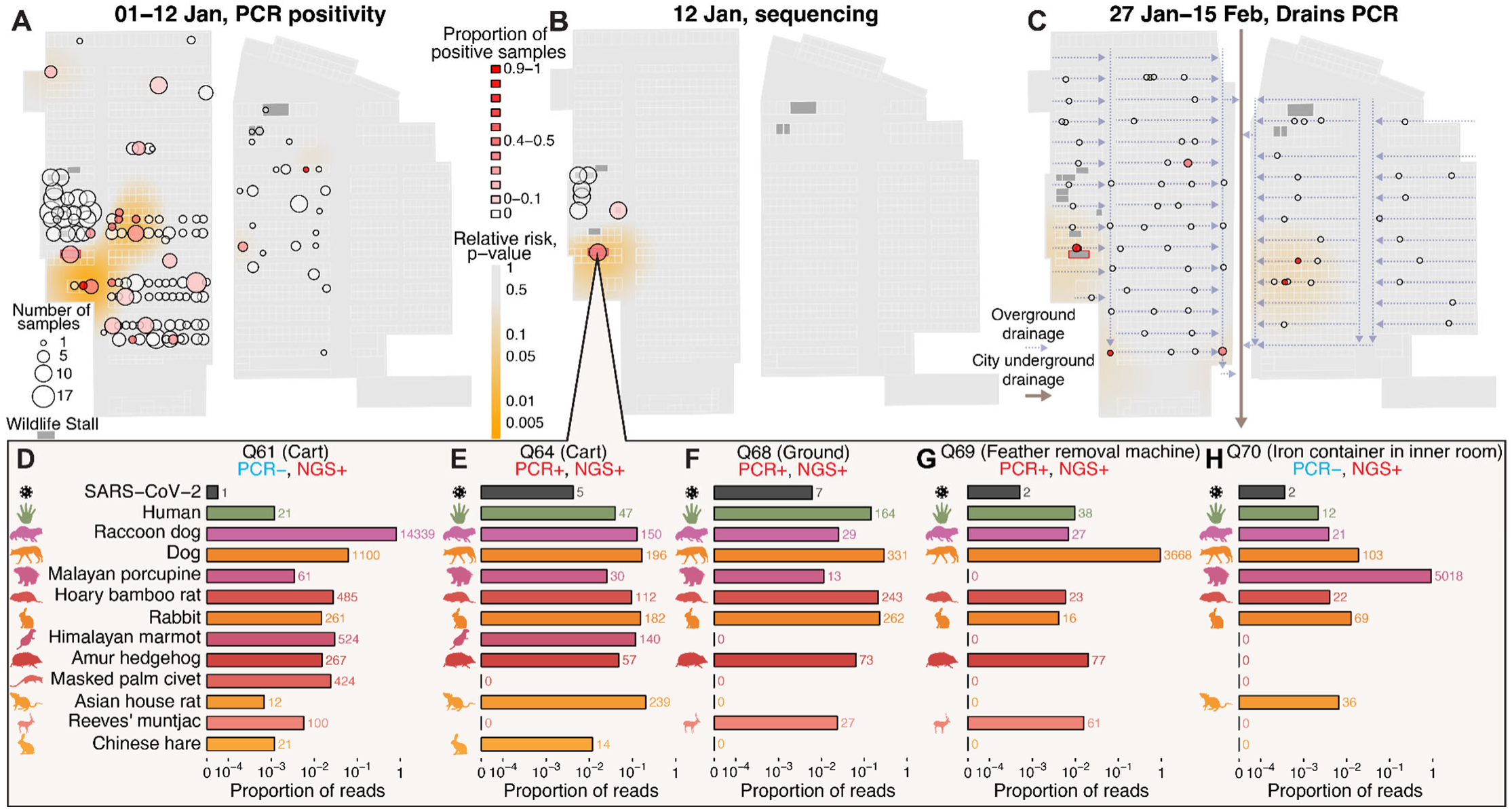
The spatial distribution of SARS-CoV-2 in the Huanan market and animal DNA/RNA in SARS-CoV-2 positive samples from wildlife stall A. **(A)** SARS-CoV-2 qPCR testing across all sampled stalls, with the proportion of positive qPCR results shown for samples collected on January 1^st^ and 12^th^ 2020. For **A–C**, the heatmaps represent the p-value surface distributions of a relative risk analysis, showing areas of significantly elevated positive test density. (**B**) Metatranscriptomic sequencing (mNGS)-based testing for SARS-CoV-2 in samples collected on January 12^th^ 2020. (**C**) SARS-CoV-2 qPCR testing of water drains tested throughout the market. Arrows indicate the direction of reported drainage flows (Liu et al. 2023). (**D–H**) Number of SARS-CoV-2 sequence reads and mammalian mtDNA reads in samples that tested positive for SARS-CoV-2 from one wildlife stall. The number after each bar is the raw number of reads. Only mammalian species reported in at least 2 samples or with greater than 300 total reads are shown.

Later sampling provided further support for wildlife stall A as a SARS-CoV-2 hotspot within the Huanan market. Of 60 samples from the underfloor drainage system of the Huanan market that were collected on January 27^th^ and 29^th^, only four were SARS-CoV-2 qPCR-positive, including the drain directly in front of wildlife stall A (**Figure 2C**). Seventeen more drain samples were collected February 9^th^ and 15^th^, with three testing qPCR-positive: one of these was again in front of wildlife stall A and the other two were from downstream locations that may have received runoff from this same stall (**Figure 2C**). Taken together, there are three independent spatial signals that identify SARS-CoV-2 positivity associated with a specific wildlife stall (A) in a section of the market with markedly higher environmental SARS-CoV-2 positivity.

The nearby wildlife stall B was also repeatedly resampled throughout January and tested qPCR-positive multiple times. In February, the offsite warehouse associated with this stall was sampled, and 5 of 12 samples tested positive for SARS-CoV-2 by qPCR (**Supplementary Table S3**). Out of the 16 samples collected from wildlife stall B on January 1^st^ and 12^th^, SARS-CoV-2 was detected only by mNGS in the January 12^th^ freezer sample, indicating lower positivity than wildlife stall A.

Outside of these wildlife stalls, the other SARS-CoV-2-positive stalls sampled on January 1^st^ were often associated with several of the known human cases in the market (**Figure S1**). These samples most likely reflect human shedding of SARS-CoV-2 in these other locations throughout the market, which was a site of ample human-to-human transmission (Worobey et al. 2022). As time of sampling progressed after the market’s closure, there was a noticeable decrease in SARS-CoV-2 viral abundance, indicating environmental viral RNA decay throughout the market over several weeks (**Figure S2**). As most wildlife stall samples were collected 11 days after the first sampling, a reduced capacity to detect SARS-CoV-2 in wildlife stall samples on January 12^th^ would then be expected due to ongoing decay of viral RNA in the environment.

### Mammalian wildlife species detected in five SARS-CoV-2 positive samples from a wildlife stall and in other wildlife stalls

Environmental samples with viral RNA can also contain genetic evidence of the mammalian hosts that shed that virus. We stringently mapped reads to a dereplicated database of eukaryotic mitochondrial genomes to quantify the abundances of animal mitochondrial DNA (mtDNA) in each environmental sample. Five SARS-CoV-2-positive samples from wildlife stall A contained mtDNA from raccoon dogs, hoary bamboo rats (*Rhizomys pruinosus*), and European rabbits (*Oryctolagus cuniculus*). Amur hedgehog and Malayan porcupine mtDNA was present in four samples, Reeves’s muntjac (*Muntiacus reevesi*) and Himalayan marmot (*Marmota himalayana*) mtDNA in three, and one sample contained masked palm civet mtDNA (**Figure 2D; Supplementary Table S6-S7**). Of these species, raccoon dogs, rabbits, and dogs (*Canis lupus familiaris*) are documented as susceptible to SARS-CoV-2 (Freuling et al. 2020; Bosco-Lauth et al. 2020; Mykytyn et al. 2021), with raccoon dogs experimentally confirmed as capable of transmission (Freuling et al. 2020). Nearby to stall A, other SARS-CoV-2-positive samples also contained wildlife mtDNA, including a garbage cart where raccoon dog mtDNA was detected and a stall with bamboo rat mtDNA (**Supplementary Table S7-S11**). While all five positive samples from stall A contained human mtDNA, humans were not the most abundant mammalian species present in these samples (**Figure 2D**). Excluding the mitochondrial 16S and 12S rRNA regions which could be differentially impacted by any potential rRNA depletion performed on these samples did not change these results (**Supplementary Table S12**; **Supplementary Fig S3**).

Further, our results show that wildlife mtDNA detection was colocalized with the reported locations of wildlife stalls (**Figure 3A, 3C–E; Supplementary Table S8-S11**). In contrast, human mtDNA was distributed throughout the market, consistent with it being a general place of human activity (**Figure 3B**). The presence of genetic material from raccoon dogs and hoary bamboo rats was highly frequent across the wildlife stalls, constituting the two most commonly detected mammalian wildlife species (**Figure 3A, C–E**). Masked palm civets’ genetic material was more rarely detected, being present in five samples from four stalls. Some wildlife species, such as nutria (*Myocastor coypus*), red foxes (*Vulpes vulpes*) and Arctic foxes (*Vulpes lagopus*) were detected only in samples and stalls that tested negative for SARS-CoV-2. To confirm species identification, we generated *de novo* contig assemblies and performed BLAST against a custom WGS database (**Supplementary Table S13**) made from available genome assemblies of species known to be at Huanan market. The BLAST results identified human and wildlife species consistent with those described above (**Supplementary Table S14**).

**Figure 3:**
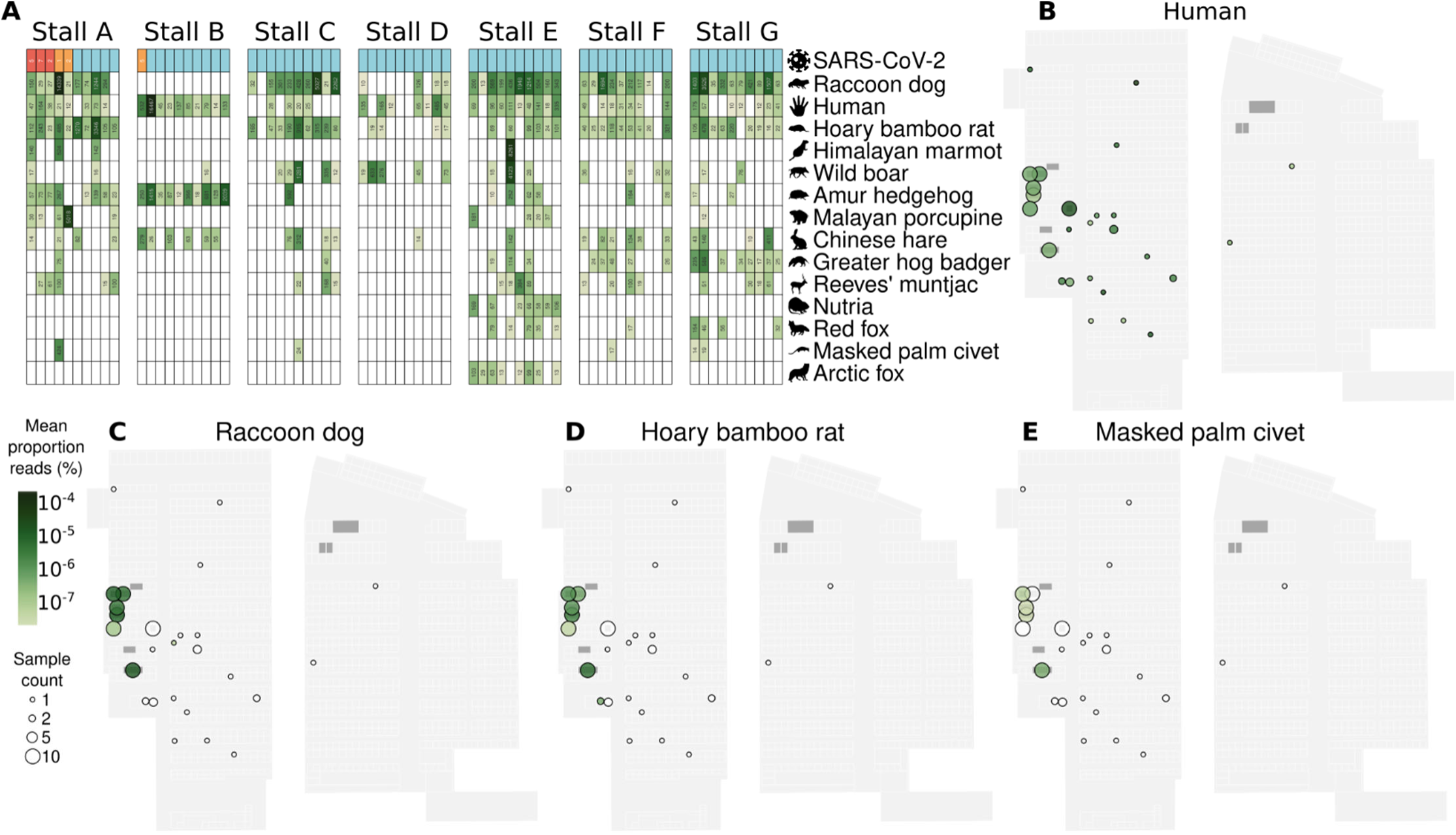
The spatial distribution of animal DNA/RNA in the Huanan market. (**A**) The sequence read counts of the mammalian species with mtDNA detected in at least 3 samples across all wildlife stall samples sequenced on January 12^th^ located in the western part of the market. Samples are grouped by wildlife stall, ordered by detection of SARS-CoV-2 (red: positive by qPCR and sequencing; orange: positive by sequencing only; blue: negative), and species are the ones we detected whose genus was reported as sold live in Wuhan markets by Xiao et al. (X. Xiao et al. 2021), ordered by abundance. (**B–E**) The spatial distribution of the mean proportions of mtDNA reads across sequenced samples collected in the market on January 1^st^ and 12^th^ for (**B**) humans (*H. sapiens),* **(C)** raccoon dogs (*N. procyonoides*), **(D)** hoary bamboo rat (*R. pruinosus)*, (**E**) masked palm civet (*P. larvata*).

Of the eighteen species reported by Xiao et. al. (Xiao et al. 2021) to be present in the four Wuhan city markets they surveyed, we confirmed genetic signatures of eleven at the species level at the Huanan market and an additional two at the genus level (**Figure 3A** and **Supplementary table S15**); Siberian weasel (*Mustela sibirica*) was rare in the market, and was absent from any of the January 12^th^ wildlife stall samples. We did not detect mtDNA sequence reads from American mink (*Neogale vison),* red squirrel *(Sciurus vulgaris),* Pallas’ squirrel *(Callosciurus erythraeus*), complex-toothed flying squirrel (*Trogopterus xanthipes*) or Asian badger in any of the environmental samples. Notably none of these, except Asian badger, was observed at the Huanan market specifically in November 2019(Worobey et al. 2022). Additionally, while Xiao et al. reported sales of Chinese bamboo rat (*R. sinensis*), we identified abundant mtDNA from the hoary bamboo rat (*R. pruinosus*) with only one sample containing trace amounts of *R. sinensis* mtDNA (**Supplementary Table S15**). This is likely due to a visual species misidentification based on physical morphology rather than genetic confirmation.

We found no evidence for the presence of any bat or pangolin genetic material, the two known hosts of sarbecovirus relatives of SARS-CoV-2, in the Huanan market. In contrast, the presence of mtDNA of *Myotis* bats was previously reported (Liu et al. 2023). To check this, we replicated this methodology by mapping reads to the Barcode of Life Data System COX1 gene database, which identified only 8 reads that mapped to any *Myotis* sequence with no mismatches. A BLAST analysis confirmed that all were non-specific matches and therefore uninformative. This indicates that neither live bats nor pangolins are likely to have been present in the sampled stalls of the Huanan market during the time period relevant to the emergence of SARS-CoV-2.

Prior studies calculated correlations of SARS-CoV-2 detection and animal sequence read abundances in market samples, concluding that SARS-CoV-2 was negatively correlated with mammalian wildlife species (Liu et al. 2023; Bloom 2021). Conceptually, these approaches are challenged by the consideration that most animal viral shedding would precede human viral shedding in a zoonotic scenario, and that most wildlife stalls were sampled 11 days after stalls with suspected COVID-19 cases (Liu et al. 2023). As a result, environmental SARS-CoV-2 RNA from non-human hosts would have had more time to decay. In addition, SARS-CoV-2 detected in non-wildlife stalls was very likely shed by humans, rendering univariate correlations including these samples inappropriate. Experimentally, the overall sampling scheme of the market sampling was also imbalanced. All wildlife stall samples from January 12^th^ were sequenced regardless of their qPCR positivity, while other sequenced samples were predominantly qPCR positive and from elsewhere in the market (**Supplementary Table S3**). Therefore, species present in wildlife stalls are disproportionately overrepresented in the sequenced negative set, and this sampling design will cause wildlife species to artifactually appear negatively correlated with SARS-CoV-2 (**Figure S4**).

As all wildlife stall samples collected on January 12^th^ had been sequenced regardless of their SARS-CoV-2 positivity, we conducted a correlational analysis of relative species abundances in these samples (*n*=70) as this could represent a balanced dataset for informing which host had shed the virus detected therein. Across these wildlife stall samples, there was no significant correlation between human mtDNA and SARS-CoV-2 RNA (ρ=0.13; 95% confidence interval [CI] [-0.09,0.34]), similar to the average mammal (ρ=0.08; 95% CI [-0.12,0.29]) (**Figure S4**). Sequence read abundances of Malayan porcupine (ρ=0.45; P<0.001, false discovery rate (FDR)=5%) and Himalayan marmot were significantly correlated with SARS-CoV-2 after multiple hypothesis correction (ρ=0.34; P<0.033, FDR=5%) (**Figure S4; Supplementary Table S16**), reflecting their increased detection in wildlife stall A. Generally, temporal trends and compositional effects in metagenomic sequencing data also influence correlations, further challenging their interpretation (Carr et al. 2019). As previously described (Crits-Christoph et al. 2023), a correlational analysis would be unlikely to provide reliable insights into whether any particular species was or was not infected by SARS-CoV-2 within the market.

### Wildlife stalls and SARS-CoV-2 positive samples contain other mammalian viruses associated with the animal trade

The presence of animal viruses with predictable host ranges provides evidence of animals productively infected with viruses in the Huanan market at the end of 2019. By mapping sequencing reads to a custom database of human and animal viruses with stringent filtering, we identified several mammalian viruses present in the market (**Supplementary Table S17**). Human-specific viruses were rare, even at a threshold of one read per sample. We found human coronavirus 229E in one stall and human respiratory syncytial virus (subgroup B) in another. Other detectable human viruses were dsDNA viruses such as human polyomavirus 6, human papillomaviruses, and human herpesviruses (**Supplementary Table S17**).

We also detected several other mammalian viruses within the market (**Supplementary Table S17**). In SARS-CoV-2 positive wildlife stalls, we identified close relatives of viruses reported to infect the wildlife species also detected in these samples (**Figure S5**). Of these viruses, close relatives of a raccoon dog amdoparvovirus, a bamboo rat betacoronavirus, and a civet kobuvirus were sufficiently abundant to reconstruct mostly complete genome sequences from samples across the market via a mapping-consensus approach. All three viruses were predominantly found in wildlife stalls with mNGS evidence of their putative hosts, and in some cases they were identified in samples from nearby locations as well (**Figure 4A–C**).

**Figure 4:**
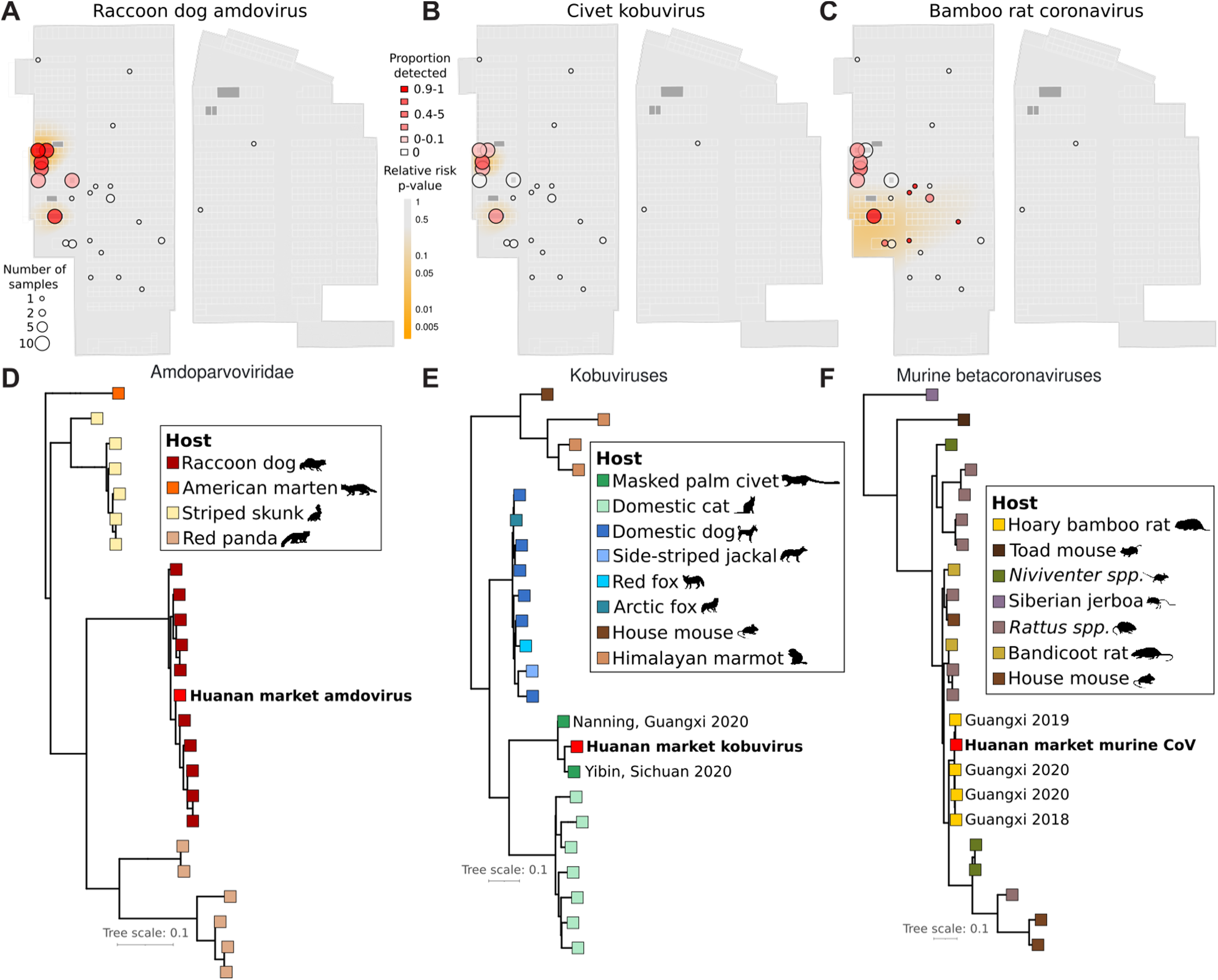
Animal viruses in the Huanan market. (**A–C**) The spatial distribution of detection of three wildlife viruses detected in the Huanan market in sequenced samples collected in the market on January 1^st^ and 12^th^. Bubbles correspond to stalls, and the bubble color represents the mean proportion of reads mapping to the viral genome from samples in that stall. The heatmap shown is a quantification of the p-value distribution for a relative risk analysis, showing spatial distribution of enriched positivity for each virus. (**D–F**) Maximum likelihood phylogenies of the market consensus sequence for each of the three viruses shown in (**A–C**). Each reference virus is colored by the species it was reported as isolated from.

We inferred maximum likelihood phylogenetic trees incorporating these viruses and their known relatives (**Figure 4D–F**). In each case, the virus present in the market was most closely related to reported viruses shed by a singular host, indicating a predictable host specificity. The two closest relatives of the civet kobuvirus we detected were from sequences identified in samples from market animals from Sichuan and Guangxi provinces, and the bamboo rat betacoronavirus was a close and recent relative of a virus identified in bamboo rats on a Guangxi farm in 2019 (Cui et al. 2023). These viruses suggest some movement of infected animals from southern China to Wuhan, a trade conduit that could have also led to the emergence of SARS-CoV-2. This result is also consistent with reports that Huanan market vendors sourced bamboo rats from Guangxi and Yunnan provinces (World Health Organization 2021). Movement of animal viruses such as these via the wildlife trade recapitulates the dispersal of SARS-CoV-1 from Yunnan to Guangdong and Hubei provinces (Shi and Hu 2008).

Additional viruses found at the market included polyomaviruses, hedgehog coronavirus HKU31, and skunk adenovirus PB1 (which has a broad host range) (**Supplementary Table S17**). Five genome segments of influenza A virus (PB2, PB1, NP, HA, and PA) were detected together in a sample from one SARS-CoV-2 negative wildlife stall. The most closely related BLAST hits of these highly fragmented segments were to H9N2 strains reported from chickens in southern China in 2017. Spillover of avian H9N2 had been reported in civets in another recent survey of market animals in China (He et al. 2022). Human zoonotic cases of H9N2 have occurred (Butt et al. 2005); hence, SARS-CoV-2 was not the only virus with zoonotic potential present in the Huanan market at the end of 2019. These results further add to the evidence for the presence of live animals at the market, and establish it as a place where potential wildlife hosts of SARS-CoV-2 were actively shedding other viruses.

### Reconstruction of mitochondrial genotypes of potential intermediate host species of SARS-CoV-2 within the Huanan market

Genotypic differences within species can be valuable for identification of the subspecies and the geographic origin of individual animals present at the Huanan market. To facilitate subspecies identification and ascertainment of the likely geographic origin of animals in the Huanan market, we reconstructed mtDNA consensus genomes from wildlife stall samples. We used a reference-guided mapping approach to obtain partial to near-complete mitochondrial consensus genomes of several mammalian wildlife species. We obtained 33 consensus mitochondrial haplotypes from separate swabs that were >50% complete compared to the reference for seven abundant wildlife species: raccoon dog, masked palm civet, hoary bamboo rat, Amur hedgehog, Malayan porcupine, greater hog badger (*Arctonyx collaris*), and the Himalayan marmot (**Figure 5A; Supplementary Table S18**). Ten of these mitochondrial genomes were >90% complete. We further identified consensus SNVs for these genomes and found they diverged on average 0.57% from the reference genome for each species (minimum: 0.16%, maximum: 2.1%).

**Figure 5:**
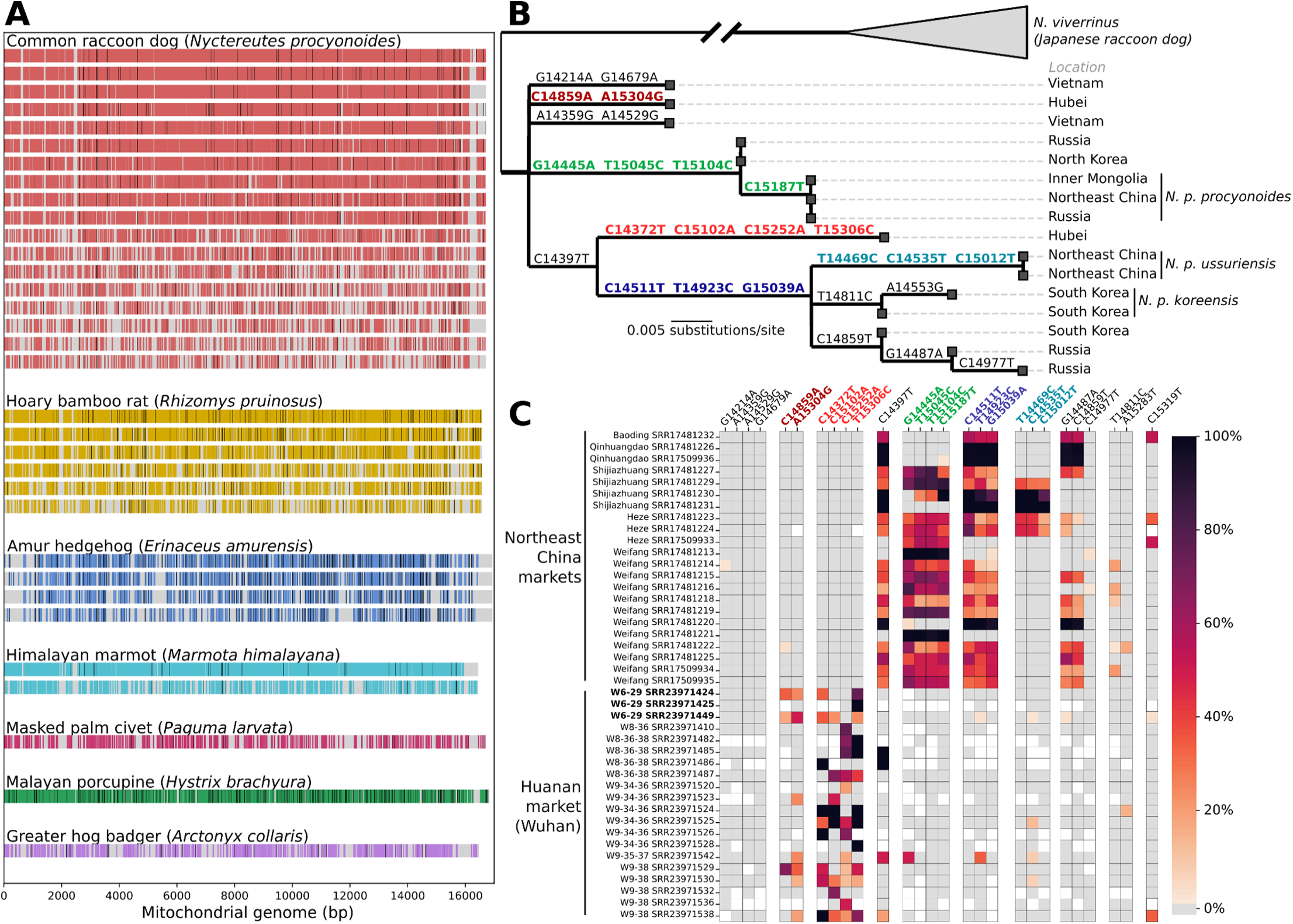
Mitochondrial phylogenetics of potential intermediate host species of SARS-CoV-2 in the Huanan market. **(A)** Coverage of wildlife mitochondrial genomes in Huanan market samples. Covered bases are colored, and consensus SNPs from the reference genome for each species are shown as black lines. (**B**) Cytochrome B phylogeny of raccoon dog reference sequences collected from mainland Asia. (**C**) Heatmap of raccoon dog *cytB* SNVs found in the Huanan market samples and samples collected from other markets to the north of Hubei province. The color of each square represents the read mapping frequency of that allele in the sample. SNVs are grouped by the branch of the reference tree they fall on, corresponding to the colors from (B). Samples from the wildlife stall positive for SARS-CoV-2 are in bold.

To investigate the geographic origins of the raccoon dogs in the Huanan market within the context of the animal trade, we analyzed their mtDNA genotypes. In Asia, the raccoon dog’s current range extends from Vietnam to Russia (Kim et al. 2013) and four subspecies are recognized: (a) *N. p. procyonoides*, found throughout China; (b) *N. p. koreensis* in the Korean peninsula, (c) *N. p. orestes* found in southern China and Vietnam; and (d) *N. p. ussuriensis*, a subspecies found in northeastern China and Russia and farmed in large numbers in this region for its thick fur (“China’s Fur Trade and Its Position in the Global Fur Industry” 2023; Shi and Hu 2008). Given the natural range of the closest known bat sarbecovirus relatives of SARS-CoV-2 in southern China, northern Laos, and Vietnam (Pekar et al. 2023; Temmam et al. 2022; Zhou et al. 2020), raccoon dogs from northern China would be an unlikely conduit for SARS-CoV-2 into Wuhan.

We performed phylogenetic inference on the mitochondrial *cytB* gene, as reference genes have been reported for wild raccoon dogs in Hubei province (2020) (Wang et al. 2022) and Vietnam (2013) (Kim et al. 2013), and multiple raccoon dog subpopulations can be distinguished using the *cytB* gene (**Figure 5B**). Because market environmental swabs can contain DNA shed by unrelated but cohoused animals of the same species, we used a metagenomic single nucleotide variant (SNV)-based approach instead of relying solely on the consensus genomes. To compare animals in the Huanan market to those from other markets, we identified SNVs for mitochondrial genomes from an independent set of samples of pooled raccoon dog samples collected from markets in five cities to the north of Hubei province during 2020 (He et al. 2022). Notably, raccoon dog samples from the other markets to the north of Hubei province were characterized by the presence of SNVs on two branches to *N. p. ussuriensis* and *N. p. procyonoides* reported from Inner Mongolia in 2016 (**Figure 5C**). These SNVs were absent from the Huanan market; instead, SNVs from two genetically distinct raccoon dog populations sampled in 2020 collected in the wild from Hubei in 2020 were present (**Figure 5C**). SNVs associated with two raccoon dog haplotypes collected in Vietnam in 2013 were absent from all samples. This result suggests that the detected raccoon dogs in the Huanan market in late 2019 were not *N. p. ussuriensis* and were a distinct population from those sold in markets in cities and raised on fur farms in northern China. It is unknown how far south the wild or farmed range of the subpopulations detected here extends due to a lack of genetic data for *N. procyonoides* in southern China. These data are consistent with a geographic origin of the raccoon dogs in the Huanan market in central or southern China, from which a viral transmission chain within the animal trade could have arisen after a spillover from a bat reservoir south of Wuhan.

## Discussion

Extensive epidemiological evidence supports wildlife trade at the Huanan market as the most likely conduit for the COVID-19 pandemic’s origin (Worobey et al. 2022; Holmes et al. 2021). While the identity of an intermediate host between the *Rhinolophus spp.* (horseshoe bat) reservoir of SARS-CoV-2-like coronaviruses and humans remains unknown, our analysis informs this open question by determining the mammalian species present in the market with species and subspecies resolution. These results show that multiple likely intermediate hosts of SARS-CoV-2 were present at the exact site within Wuhan at which COVID-19 was first epidemiologically linked. It is not possible to conclude which of these species may have been infected and/or introduced the virus to the market from this data alone. Nonetheless, our analysis provides a small and actionable list of species with genotypic details. Of the wildlife species detected in SARS-CoV-2 positive environmental samples, four have previously been implicated in bat coronavirus cross-species transmission through the animal trade: raccoon dogs, masked palm civets, hoary bamboo rats, and Malayan porcupines (Guan et al. 2003; Huong et al. 2020).

Among the potential intermediate hosts present in the Huanan market, raccoon dogs are known to be susceptible to SARS-CoV-2, to shed high titers of virus, and to be able to transmit (Freuling et al. 2020). The common raccoon dog was the most abundantly detected animal species in market wildlife stalls sampled on January 12^th^, and in the wildlife stall with the most SARS-CoV-2 positive samples (**Figure 3A; Supplementary Table S15**). The experimental susceptibility of civets is unknown, but *Paguma larvata* cells are susceptible to pseudotyped VSVs expressing the SARS-CoV-2 Spike protein *in vitro* (Li et al. 2023). However, the susceptibility of bamboo rats, Malayan porcupines, and Amur hedgehogs remains unknown, and these species should also be prioritized for susceptibility testing. Of the other species in the market, marmots may be an unlikely conduit for SARS-CoV-2 considering that their usual range is at very high elevations (Wu et al. 2023), and muntjac deer have a truncated ACE2 gene without a signal peptide sequence that suggests a lack of susceptibility (GenBank accession: NP_001358344.1). Siberian weasels (Boklund et al. 2021; Shi et al. 2020), foxes (Porter et al. 2022), and greater hog badgers (Davoust et al. 2022) are either known or closely related to species thought to be susceptible to SARS-CoV-2, although these species were very rare or absent in SARS-CoV-2 positive samples and stalls. Other market mammals are known to be susceptible to SARS-CoV-2 but do not represent a significant transmission risk, including dogs, rabbits, and boar, as infected animals do not produce or shed virus at high titers (Bosco-Lauth et al. 2020; Mykytyn et al. 2021; Meekins et al. 2020; Bosco-Lauth et al. 2021).

Multiple lines of evidence are consistent with the infection of wildlife animals with SARS-CoV-2 in the Huanan market. Animal carts, a cage, and a hair/feather remover from a wildlife stall tested positive for SARS-CoV-2, and there was more DNA from mammalian wildlife species in these samples than human DNA. The surrounding stalls also had relatively higher rates of SARS-CoV-2 positivity, and drains adjacent to and downstream of this wildlife stall tested positive for SARS-CoV-2. Finally, there were several other viruses known to infect wildlife in these samples. These data indicate either that the animals present at this stall shed the SARS-CoV-2 detected on the animal equipment, or that early unreported human case(s) of COVID-19 shed virus in the exact same location as the detected animals. While either or both scenarios are consistent with these data, only a zoonotic origin of SARS-CoV-2 directly predicts co-detection of SARS-CoV-2 and wildlife genetic material.

It has been proposed that humans could have introduced the virus into the Huanan market (Liu et al. 2023; Bloom 2023). It is most likely that there were human infections of SARS-CoV-2 earlier than the first documented and hospitalized market cases, including unascertained market cases or contacts thereof. However, the detection of both lineage B and lineage A within and indirectly linked to the Huanan market implies that SARS-CoV-2 most likely emerged there. Any hypothesis of COVID-19’s emergence has to explain how the virus arrived at a wildlife market in a city of Wuhan’s size at a time when so few humans were infected (Worobey et al. 2022). Human introductions linked to the animal trade offer one explanation for this, and the introduction of the virus by an animal trader or farmer cannot be excluded, but these hypotheses are challenged by phylodynamic evidence for multiple spillovers (Pekar et al. 2022).

The detection of SARS-CoV-2 RNA in the Huanan market in January 2020 could plausibly reflect deposition several weeks before sampling, compatible with estimated dates of the first human infections. SARS-CoV-2 RNA has been detected on indoor surfaces for prolonged periods spanning several weeks (Renninger et al. 2021; Liu et al. 2021; Zhou et al. 2023), and the temporal signal of viral RNA decay in the Huanan market samples themselves offers further support for these timescales (**Figure S2**). While the earliest zoonotic events of SARS-CoV-2 most likely occurred in late November 2019 (Pekar et al. 2022), infected cohoused animals could be expected to shed virus for weeks longer. SARS-CoV-2 detected in the Huanan market may be remnant from any time in that period of unknown length.

Focused genetic and serological sampling of raccoon dogs and the other mammalian species reported here throughout Southeast Asia and southern China can shed light on the animal trade networks that may have led to the emergence of SARS-CoV-2 as previously recommended (World Health Organization 2021). Serological testing of the oldest animals (for instance the breeding stock) in source farms might provide additional information of transient circulation, as has been observed in mink farms infected with SARS-CoV-2 (Rasmussen et al. 2021). Future studies to clarify the susceptibility status of all of these species using *in vitro* approaches and live-animal infection experiments, should also be prioritized. The limited viral and serological sampling of these species in Southeast Asia and southern China (Wang et al. 2022; Wang et al. 2022) indicates that the wildlife trade directly before the COVID-19 pandemic is highly undersampled, or underreported. Retrospective studies should be performed, where possible, testing the species described here throughout the animal supply chains of Southeast Asia and southern China, through which in all scientific likelihood the COVID-19 pandemic emerged.

## Methods

### Sample preprocessing

184 sequencing runs from the NCBI BioProject PRJNA948658 were downloaded and quality trimmed using BBDuk using the settings: ktrim=rl k=17 qtrim=r trimq=10 maq=10 minlen=30 entropy=0.5 threads=60 ref=./all_adapters.fa. A FASTA file of adapters which included BGI adapters was passed for trimming.

### SARS-CoV-2 mapping

Reads were mapped to the SARS-CoV-2 reference genome (NC_045512.2) using Bowtie2 (Langmead and Salzberg 2012) and default settings. Mapped reads were filtered to those with at least 97% identity to the reference, a minimum mapping quality score of 20, a minimum alignment length of 95%, and reads mapping at least 200 bp away from the contig edge, and counted using a custom Python script (“count_reads_sars2.py”). For paired read samples, a mapped read pair was counted as a single observation.

### SARS-CoV-2 phylogenetics

ViralConsensus v0.0.3 (Moshiri 2023) was run to generate consensus genomes and examine SNVs for the samples with the settings: --min_qual 20, -- min_depth 10, --min_freq 0.5, --ambig N. The iVar pipeline (Grubaugh et al. 2019) was run to generate consensus genomes and examine SNVs for the A20 sample, as this sample was generated with an amplicon based approach. 15 bp were trimmed from the 5’ and 3’ ends of reads and a minimum depth of coverage of 15x was required. We augmented the data set of SARS-CoV-2 genomes from Pekar et al. 2022 (J. E. Pekar et al. 2022) (those collected by Feb 14, 2020) using the four reconstructed SARS-CoV-2 genomes from the environmental samples (the two SARS-CoV-2 genomes from Huanan market environmental samples present in the dataset from Pekar et al. were excluded, as they were derived from 2 out of 4 environmental samples here). Molecular clock phylodynamic inference was conducted using a Bayesian approach in BEAST v1.10.5 following the same protocol as in Pekar et al. We employed a non-reversible, random-effects substitution model, a strict molecular clock, and a non-parametric skygrid prior with 20 grid points and a cut off of 0.37, which translates to 5 October 2019.

We performed three analyses: (i) an analysis using the 17 market-associated genomes (13 identified cases and the 4 genomes reconstructed from the environmental samples), (ii) a Wuhan-focused analysis using the 93 genomes from Wuhan, and (iii) an analysis using 789 genomes, representing the early global diversity of SARS-CoV-2. For (i), we ran one chain of 100 million generations, subsampling every 10 thousand generations to continuous parameter log files and the tree file. For (ii), we ran one chain of 100 million generations, subsampling every 100 thousand generations to continuous parameter log files and the tree file. For (iii), we ran three independent chains of 400 million generations, sub-sampling every 25 thousand iterations to continuous parameter log files and 100 thousand iterations for the tree files. The first 10% of each chain was discarded as burn-in. Convergence and mixing was assessed in Tracer v1.7.1, and the 3 chains for analysis (iii) were combined in LogCombiner. All relevant effective sample size (ESS) values were >200 for the final log file for each analysis. The accession IDs can be found in **Supplementary Table S19**.

### Mitochondrial mapping

All eukaryotic mitochondrial genomes were downloaded from NCBI’s RefSeq and GenBank databases to build a custom mapping mitochondrial database. GenBank sequences with “partial”, “gene”, “genome assembly”, and “chromosome” in the description and those smaller than 12 Kb were removed. Genomes were clustered by Mash (Ondov et al. 2016) distances, first at 98% identity, preferentially selecting RefSeq genomes as cluster representatives. Reads were mapped with Bowtie2 to the 98% identity genome index and mapped reads with >=95% identity, MAPQ>=20, and mapping lengths >= 40 were retained using a custom Python script (count_reads98.py). For paired read samples, a mapped read pair was counted as a single observation. Next, a second round of reference genome clustering at 93% identity was performed, preferentially selecting the cluster representative as the genome with the highest sum of covered bases across all market samples. Reads were mapped again with Bowtie2 to this ‘93% clustered genome index’ and counted using a custom Python script (count_reads93.py) and similar cutoffs as described above. The resulting hits were filtered to Metazoa and assigned taxonomy with the Taxoniq package. To assess the potential differential impact of rRNA depletion on different species, we queried the mitochondrial genomic positions of the 16S and 12S rRNAs for all mammalian species observed in the market. A custom Python script was used to count mapped reads filtered in the same way as above, except also excluding all reads that overlapped with the genomic positions of the 16S or 12S for each mitochondrial genome.

### Mapping and analysis of environmental samples

We enhanced the market geospatial map from Worobey et al. (Worobey et al. 2022) using data on environmental samples taken from the market(Liu et al. 2023), including both SARS-CoV-2 positive and negative samples. We could precisely locate 783 of the 819 samples from inside the Huanan market. The resulting sample locations and associated metadata were integrated with reported qPCR results and the results of our mNGS mapping for downstream analyses. Samples were grouped by stall to calculate the fraction of positive samples or the average proportion of reads associated with the species of interest. Overground and city drainage paths were plotted in accordance with published drainage maps(Liu et al. 2023).

### Spatial relative risk analyses of environmental samples

As in Worobey et al.(Worobey et al. 2022), spatial relative risk analyses were performed for SARS-CoV-2 and other key viruses using the “sparr” package available in R (Davies, Marshall, and Hazelton 2018), with linear boundary kernels for edge correction and bandwidth selection using least-squares cross validation. For analyses including market drains, we used a wider market boundary that included the drain sites outside of the market building. We studied variation in the relative risk quantity r(z)=f(z)/g(z) at each position z, where f(z) is the test distribution and g(z) is the control distribution, and tested the null hypothesis H_0_: r(z) = 1, against the alternative hypothesis of increased relative risk, H_1_: r(z) > 1. We then plotted an asymptotic p-value approximation P(z), a pointwise estimate of statistical significance.

### Correlational analyses

Spearman’s rank correlation coefficients and p-values were calculated using the scipy package between animal species abundances (mitochondrial mapping results) and SARS-CoV-2 mapped read counts, both normalized to total number of reads per sample after pre-processing. Reported conclusions were robust against normalization method (total reads, mapped reads, or no normalization). Species were included in analysis if they were identified in three or more samples. Reported correlation coefficients and 95% CI were estimated by bootstrapping with 1000 permutations. P-values for statistical significance of correlations of mammalian species and SARS-CoV-2 in wildlife stall data were corrected for multiple hypotheses using the Benjamini/Hochberg procedure implemented in the statsmodel package.

### Viral mapping and assembly

A viral database of all viruses deposited in the RefSeq database was created and supplemented with viral genomes from two recently published studies (Cui et al. 2023; He et al. 2022) that reported viruses from wildlife animals. This set of genomes was clustered by Mash distances at >95% nucleotide identity. Low-complexity regions of these viruses were detected using dustmasker 1.0.0 with default settings. Reads from each sample were mapped to this dereplicated viral database using Bowtie2. The resulting mappings were filtered with a custom Python script (count_reads_viral.py) that counted reads and/or read pairs and covered bases of each viral genome with the following filters: mapping quality >30, read alignment length >95%, read percent identity to the reference >97%, and base pairs mapping to within 200 bp of the contig edges, or to low-complexity regions, were ignored. Only viral genomes with a breadth of coverage of at least 500 genomic nucleotides in at least one market sample were retained.

To assemble consensus genomes for the raccoon dog amdovirus, bamboo rat coronavirus, and civet kobuvirus, we used a reference-guided co-assembly approach due to low sequencing coverages of viruses in the data using a custom Python script (get_viral_consensus.py). Reads from all samples with at least 500 bp of genomic breadth of coverage of the most closely related reference genome were pooled, and the consensus genotype of all mapped reads at each genomic position was used to infer the consensus genomes. Reference positions with genotype ties or no mapped reads were filled with ‘N’. To identify Influenza H9N2 partial genomic fragments, we first noticed reads mapped to multiple Influenza A genome fragments in sample SRR23971532. Because our viral read mapping based approach does not distinguish closely related segmented virus subtypes, we reconstructed the partial consensus sequences for these segments from the sample and performed a BLAST against the NR database. The PB2 sequences, for which we had the best coverage, were more closely related to H9N2 sequences; other fragments hit H9N2 and H7N9 equally well. However, three reads mapping to the HA protein had BLASTN 100% identity to the H9N2 HA gene, confirming the PB2 placement of these sequences as H9N2. The codetection of genome segments in the same sample greatly increased confidence in this call.

### Viral phylogenetics

We collated viral genome sequences from Genbank (299 amdoparvovirus, 283 kobuvirus) and aligned them with the Huanan Market sequences using MAFFT v.7.490 (Katoh and Standley 2013) with default parameters. For amdoviruses we proceeded with phylogenetic inference using the full genome alignment, but downsampled to the RdRp-encoding region of the kobuvirus RNA genome. We inferred a maximum likelihood tree with IQ-TREE 2 v2.0.7(Minh et al. 2020), using a GTR+F+G4 model with 1000 bootstrap replicates. The trees were midpoint rooted.

We aligned the Huanan Market bamboo rat coronavirus sequence with 54 *Embecovirus* full genomes from NCBI using MAFFT v.7.490 with default parameters. Because the market sequence is fragmented, we removed all regions from the alignment where it consisted of Ns, leaving a concatenated alignment of 29,468 nucleotides. We used this alignment to infer a maximum likelihood tree with IQ-TREE 2 using a GTR+F+G4 model with 1000 bootstrap replicates. We midpoint rooted the tree for analysis and visualization.

### Mitochondrial genotype reconstruction

A reference-guided mapping based approach was used to reconstruct mitochondrial consensus genomes from each sample. Reads were mapped to the eukaryotic mitochondrial database as described above, and for mammalian wildlife species, the consensus base at each position was used to infer the consensus genome with a custom Python script (mt_consensus_genomes.py), filling in all reference positions without coverage with ‘N’. Mitochondrial genomes are shared via the GitHub repository for this work.

### Cytochrome B phylogenetics

We collected 44 published raccoon dog mitochondrial sequences and aligned them to the reference (NC_013700.1) using MAFFT (options --auto --keeplength -- addfragments). As most of these were only of the cytochrome B (*cytB*) gene, we performed phylogenetic inference using only the cytochrome B gene. We removed two genomes with haplotypes identical to other genomes from Kim et al. (2013) and then inferred a maximum likelihood tree with IQ-TREE 2 v.2.0.7. using a generalized time reversible model with four gamma rates (GTR+G4). The tree was midpoint-rooted, and we then used TreeTime v0.8.1(Sagulenko, Puller, and Neher 2018) to perform ancestral sequence reconstruction. Trees were visualized using baltic 0.2.2 (https://github.com/evogytis/baltic).

We inferred a maximum likelihood tree using the entire mitochondrial genomes (genomes where only the cytochrome B gene was available were padded with Ns) with IQ-TREE 2. The tree was midpoint-rooted, and then we used TreeTime to perform ancestral sequence reconstruction. The inferred sequence for the most recent common ancestor (MRCA) of the non-Japanese clade of raccoon dog genomes (Fig. 5b) was used as a reference genome for subsequent analyses. Using pysam 0.21.0, we calculated the major allele frequency, minor allele frequency, allele frequency matching the reconstructed MRCA sequence, and allele frequency matching the inferred cytochrome B substitutions relative to the reconstructed MRCA sequence (Fig. 5b). We created heatmaps with the latter mutation allele frequencies using Seaborn 0.12.2.

### Transcriptomics assembly and BLAST

The *de novo* transcriptomic assemblies for 180 adapter cleaned samples were generated using rnaSPADES (v3.15.5) (Bushmanova et al. 2019). The resulting assembled transcripts were searched using blastn (v 2.14.0+)(McGinnis and Madden 2004) against the NCBI non-redundant nucleotide database (last update; 29^th^ May 2023) downloaded on 1 June 2023 https://ftp.ncbi.nlm.nih.gov/blast/db/) using blastn (v 2.14.0+). The specific parameters used are: -outfmt ’6 qseqid sseqid pident evalue score bitscore length qstart qend sstart send stitle’ -max_target_seqs 2. The output files were filtered to exclude hits with alignment length of less than 100. BLAST (v2.14.0+.) searches were also performed against the in-house database of genome sequences assemblies from 108 animal species (Excel sheet with accession numbers). The genome assemblies were downloaded from the NCBI genome database. The blastn search parameters used are: -outfmt ’6 qseqid sseqid pident evalue score bitscore length qstart qend sstart send stitle’ -max_target_seqs 2. The output files were filtered to exclude hits with alignment length of less than 300 bp and <99.5% nucleotide identity to the reference.

## Data availability

Analysis scripts, genome FASTA files, BAM files, and intermediate data files are available at a GitHub repository associated with this work: https://github.com/sars-cov-2-origins/huanan-market-environment

Animal silhouettes in the figures are provided by the Phylopic R package(Gearty and Jones 2023).

## Author contributions

Conceptualization: F.D., M.W., K.G.A., A.C-C.; Formal analysis: A.C-C., F.D., J.I.L., J.E.P., S.A.G., R.S., Z.H., K.G., M.B.R.; Data curation: F.D., N.M., A.C-C., J.I.L, J.E.P., R.S.; Investigation: F.D., M.W., A.C-C., Z.H., J.I.L., R.F.G., A.R., J.O.W.; Software: K.G., N.M., R.S.; Project administration: F.D., M.W., K.G.A., A.L.R.; Resources: K.G.A., A.L.R., J.O.W.; All authors wrote and edited the paper.

## Acknowledgements

We gratefully acknowledge all data contributors for generating the data on which our analyses are based. The Huanan market sequence data were generated by Liu et al.(Liu et al. 2023) and shared via NCBI BioProject PRJNA948658. We have also analyzed data in this work from BioProject PRJNA793740 and PRJNA795267. Figure 1 includes data shared on GISAID (**Supplementary Table S19**), and we gratefully acknowledge the authors from the originating laboratories and the submitting laboratories, who generated and shared through GISAID the viral genomic sequences and metadata on which this research is based(Khare et al. 2021).

This project has been funded in part with federal funds from the National Institute of Allergy and Infectious Diseases, National Institutes of Health (NIH), Department of Health and Human Services (contract no. 75N93021C00015 to M.W.). This work was partially supported through US National Institutes Health grants U19 AI135995 (KGA, RFG, MAS), U01 AI151812 (KGA, RFG), R01 AI153044 (MAS, PL, AR), R01 AI135992 (JW), 5T32AI007244-38 (JIL), EU commission H2020 programme grant number 874735 (MPGK), Fundação para a Ciência e a Tecnologia (FCT) through MOSTMICRO-ITQB (UIDB/04612/2020, UIDP/04612/2020, ZH) and LS4FUTURE (LA/P/0087/2020, ZH), Wellcome Trust (Collaborators Award 206298/Z/17/Z, ARTIC network) (MAS, PL, AR), European Research Council (grant agreement no. 725422 – ReservoirDOCS) (MAS, PL, AR), National Institutes of Health (T15LM011271; JEP), the UC San Diego Merkin Fellowship (JEP), National Health and Medical Research Council, Australia (GNT2017197, ECH), UK Medical Research Council (MRC, MC_UU_12014/12, MC_UU_00034/5 and MR/V01157X/1, DLR), AIR@InnoHK administered by the Innovation and Technology Commission, Hong Kong Special Administrative Region, China (ECH), European Union Horizon 2020 (project MOOD, grant agreement n°874850, PL), Research Foundation - Flanders (G0D5117N, G051322N, PL), the Canadian Institutes of Health Research as part of the Coronavirus Variants Rapid Response Network (CoVaRR-Net; CIHR FRN#175622; ALR, RS), and the Canada Foundation for Innovation – Major Science Initiatives Fund and from the Government of Saskatchewan through Innovation Saskatchewan and the Ministry of Agriculture (ALR, RS, MBR)

## Supplementary Table Legends

Table S1: iVar SNVs of sample SRR23971533 (A20)

Table S2: Known early sequenced SARS-CoV-2 cases from December 2019. Table S3: All sample metadata, adapted and extended from (Liu et al. 2023).

Table S4: All sample metadata for sequencing data from BioProject PRJNA948658. Table S5: SARS-CoV-2 sequence read counts from each sample.

Table S6: Scientific and common names of animals referred to in this study.

Table S7: Mammalian mtDNA detection in SARS-CoV-2 positive samples.

Table S8: Mammalian mtDNA sequence read counts in each sample.

Table S9: Mammalian mtDNA genome breadth of coverage.

Table S10: All animal mtDNA sequence read counts.

Table S11: All animal mtDNA genome breadth of coverage.

Table S12: Mammalian mtDNA sequence read counts excluding rRNA regions.

Table S13: BLAST WGS database built for transcriptome contig taxonomic assignment.

Table S14: Closest BLAST hits of contigs in SARS-CoV-2 positive samples from wildlife stall A.

Table S15: Comparison of mammals observed at the Huanan market to mammals detected in environmental sequencing data.

Table S16: Correlations between SARS-CoV-2 and mammalian mtDNA sequence read abundances.

Table S17: Sequence read abundances and genome coverage of viruses detected in this study. Table S18: Statistics of mammalian mtDNA consensus genotypes reconstructed in this study. Table S19: GISAID accession numbers of genomes used in this study.

---

**Figure S1:**
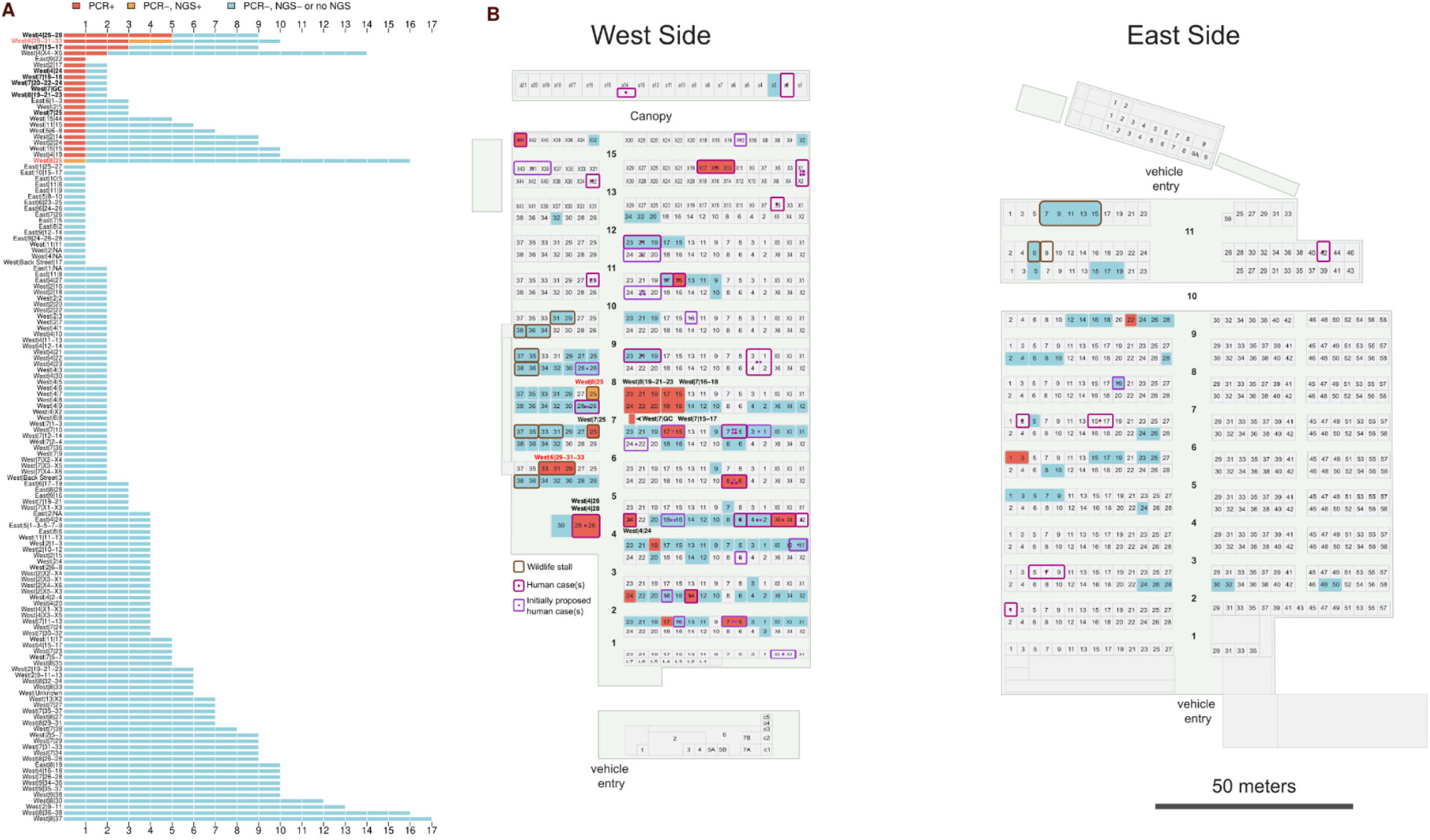
A. SARS-CoV-2 positivity by stall for samples collected on January 1^st^ and 12^th^, and B. market sampling map.

**Figure S2:**
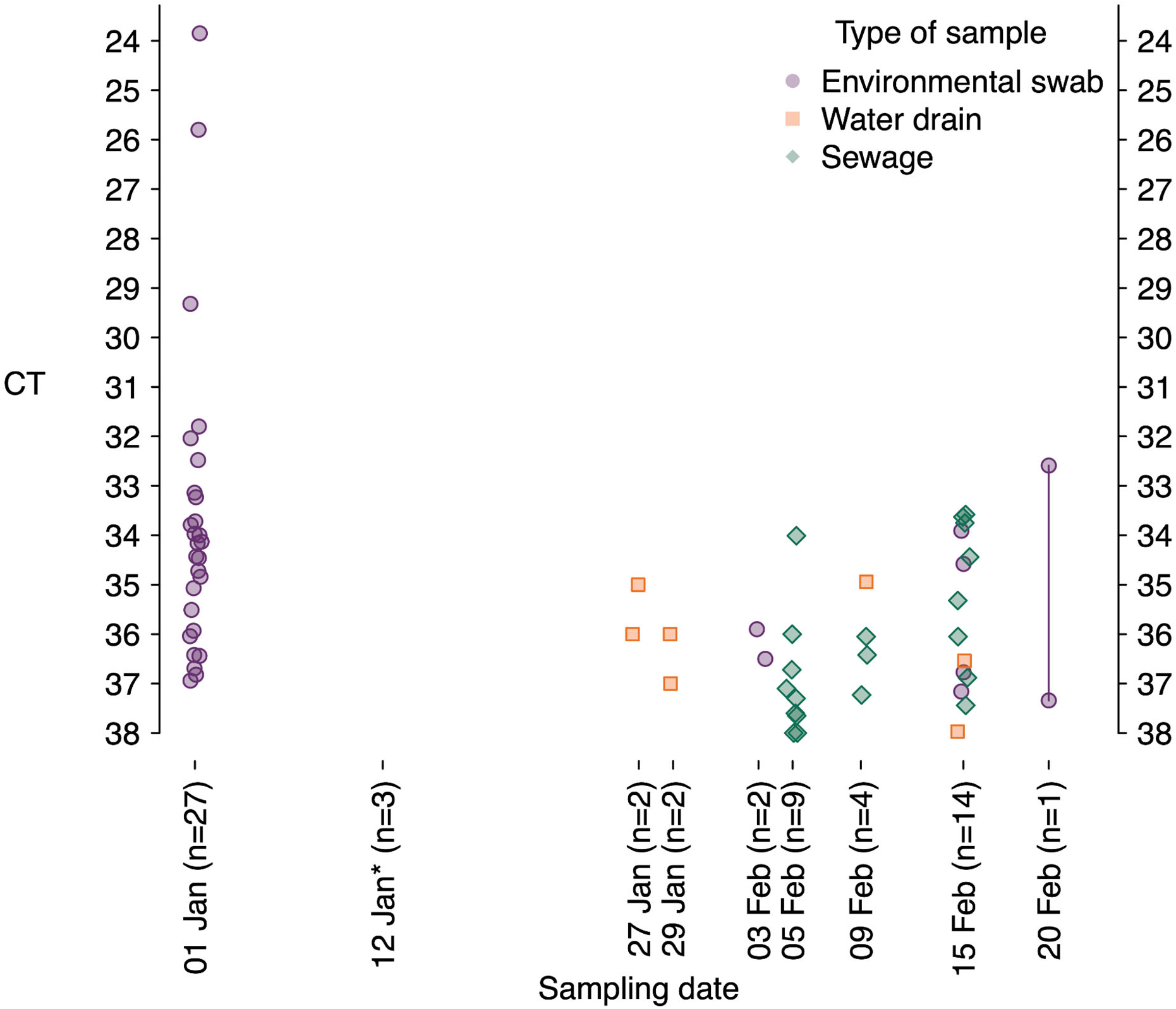
SARS-CoV-2 qPCR Ct values by date of sampling. *: Ct values were not recorded by Liu et al. for the January 12^th^ samples (3 were positive by qPCR). Two Ct values are available for the February 20^th^ sample. Data from Liu et al. Supplementary Table 2.

**Figure S3:**
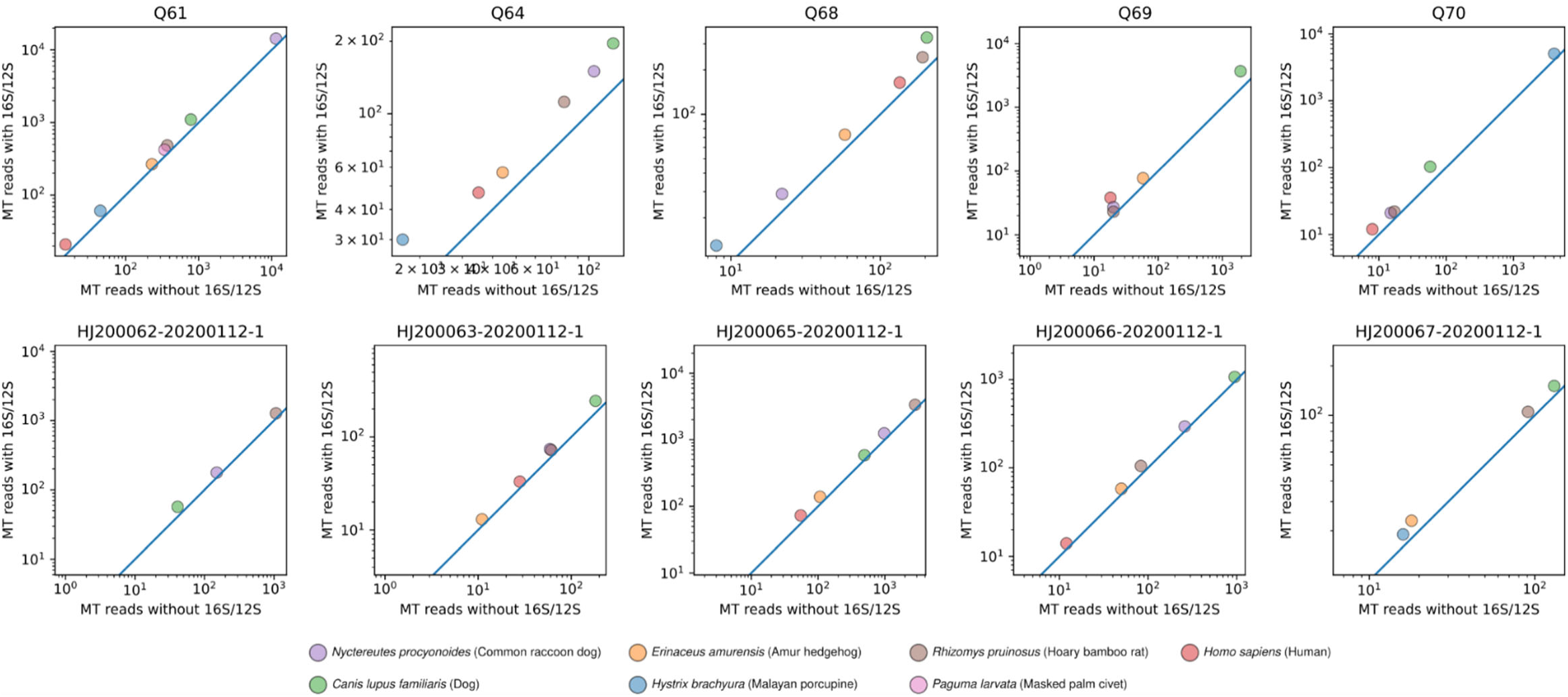
Abundances of mitochondrial DNA from humans and key wildlife species in SARS-CoV-2 positive (top row) and negative (bottom row) samples from Stall 6-29 with and without including mitochondrial rRNA regions.

**Figure S4:**
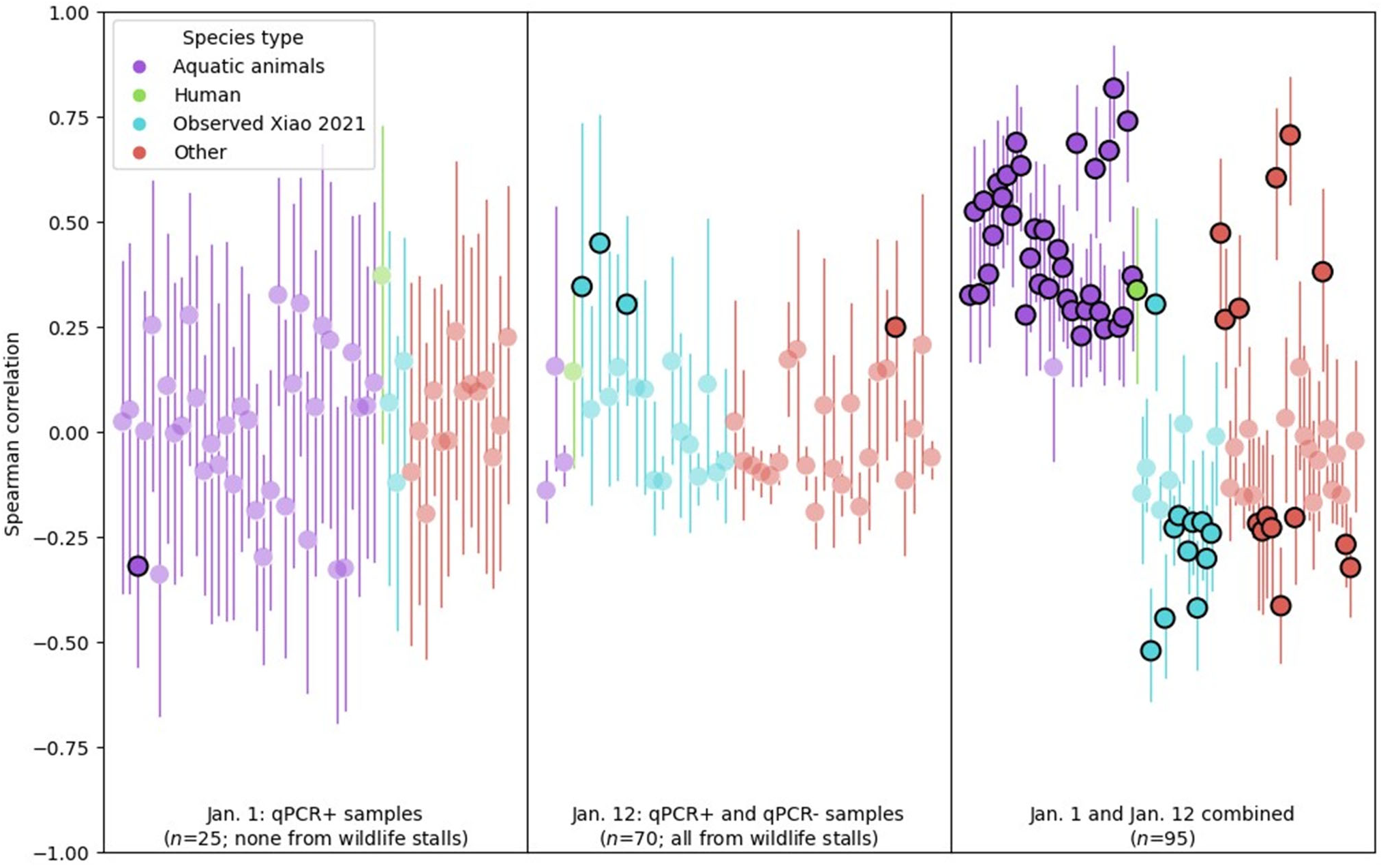
Spearman correlations between animal species abundances and SARS-CoV-2 reads in different sample sets. The estimated Spearman correlation coefficient and its 95% CI are shown for species detected in 3 or more samples collected on January 1^st^, January 12^th^, or either date. Highlighted points have uncorrected p-values below 0.05.

**Figure S5:**
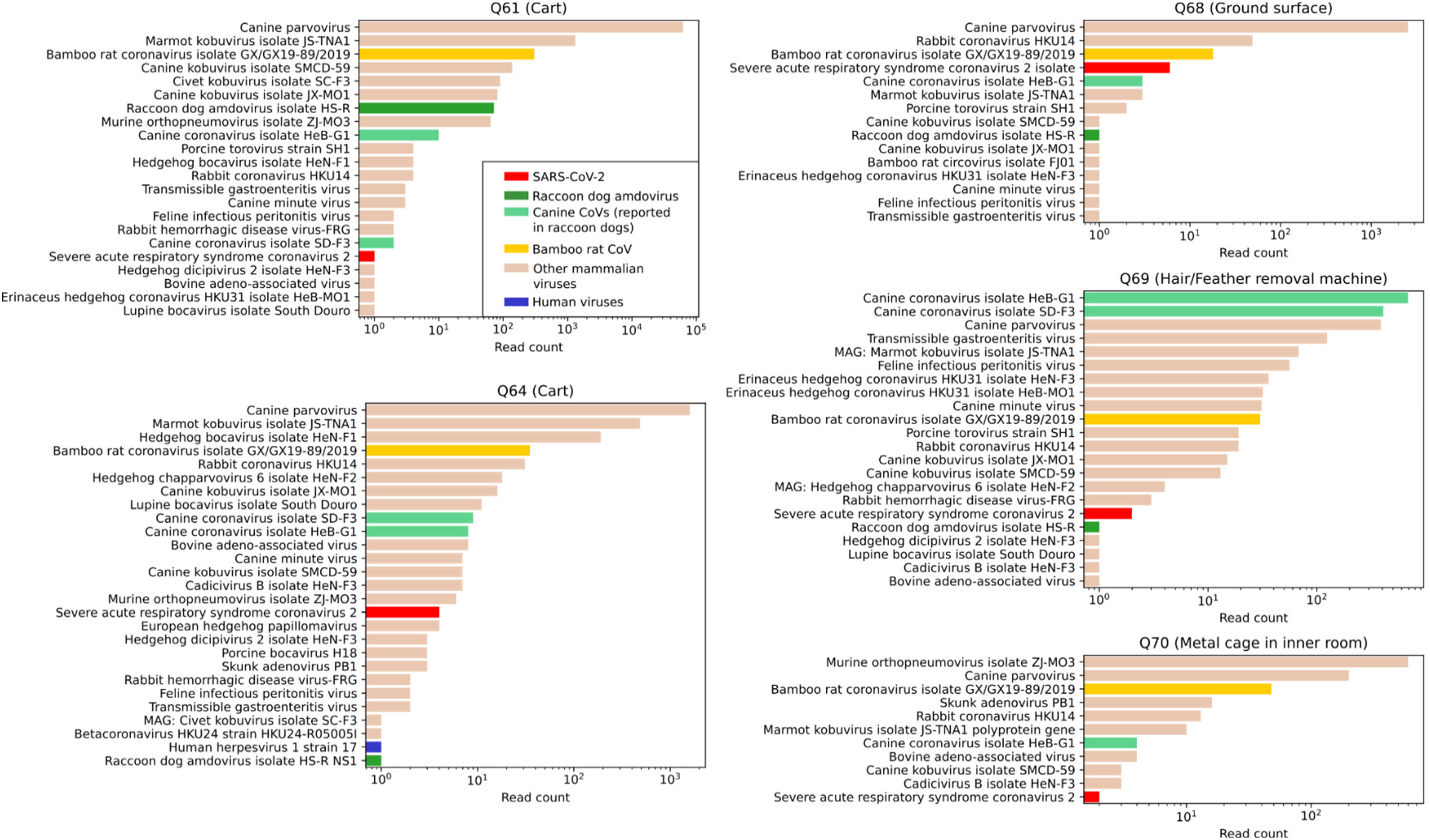
Viral abundances within 5 SARS-CoV-2 positive samples from a wildlife stall.

